# Single-cell transcriptomics of peripheral blood in the aging mouse

**DOI:** 10.1101/2021.04.08.439040

**Authors:** Yee Voan Teo, Ashley Webb, Nicola Neretti

## Abstract

Compositional and transcriptional changes in the hematopoietic system have been used as biomarkers of immunosenescence and aging. Here, we use single-cell RNA-sequencing to study the aging peripheral blood in mice, and characterize the changes in cell-type composition and transcriptional profiles associated with age. We identified 17 clusters from a total of 14,588 single cells. We detected a general upregulation of antigen processing and presentation and chemokine signaling pathways and a downregulation of genes involved in ribosome pathways with age. We also observed increased percentage of cells expressing markers of senescence, Cdkn1a and Cdkn2a, in old peripheral blood. In addition, we detected a cluster of activated T cells that are exclusively found in old blood, with lower expression of Cd28 and higher expression of Bcl2 and Cdkn2a, suggesting that the cells are senescent and resistant to apoptosis.

## Introduction

The functional decline of the immune system with age, or immunosenescence, is associated with different hematopoietic changes, including a decrease in the replication ability of hematopoietic stem cells and in B lymphopoiesis, and lower efficiency in CD4 and CD8 T cells response (1). It has been shown that with age the immune cells’ composition shifts, with a reduction of B cells in the blood in old individuals; increased B1 cells, activated and memory B cells in old peripheral blood in mice; increased levels in the bone marrow in aged mice; increased proportion of NK cells in human blood; and decreased number of NK cells in mouse blood with age (2-6). Some of these shifts have been proposed as markers for biological aging as the could predict longevity (7). T cells can undergo replicative senescence in human aging after extensive antigen-driven proliferation. These cells can be identified by the Cd8+ Cd28-markers. Donor-specific replicative senescent T cells found in organ transplant patients have been shown to be tolerable to rejection, suggesting a possible suppressive mechanism of these cells in reducing reactivity against allograft (8, 9). Several studies have observed a significant correlation between the number of Cd8+ Cd28-T cells and the decline of antibody response to influenza vaccination in the elderly (9, 10).

Aging also has profound effects on hematopoetic stem cells (HSCs) as well. The number of lymphoid HSCs decreases with age whereas the number of myeloid HSCs increases with age. Beside the quantity, the quality of lymphoid HSCs also declines with age, resulting in a lower proliferation rate and a higher apoptosis rate in T and B lineage progenitor cells (11). Beyond the decline in the immune function and in the number of cells and declining with age, the blood also contains other factors that can modulate aging. Parabiosis of young and old mice has been shown to reverse cardiac hypertrophy in old mouse and rejuvenate aged satellite cells (12, 13). Aging has also been shown to increase cellular heterogeneity (14, 15). For example, a recent study using single-cell RNA-seq (scRNA-seq) showed that transcriptional heterogeneity of CD4+ T cells increased upon stimulation in old mice compared to young mice, indicating that gene expression of immune cells is dysregulated with age (15).

Here, we applied scRNA-seq on young and old mice to dissect the transcriptional and cell composition changes of all cell types in the peripheral blood with age.

## Materials and Methods

### Use of Animals

C57BL/6 female 4 month and 24 month mice were obtained from the National Institute of Aging (NIA). They were fed ad libitum and kept in standard housing conditions and all procedures were approved by the Brown University Institutional Animal Care and Use Committees (IACUC) committee.

### Isolation of immune cells from the peripheral blood

Blood samples were drawn from the heart of four young and four old mice. Samples from two mice within each age group were pooled, resulting in two pooled young and two pooled old samples. Subsequently, the pooled samples were diluted 1:1 with PBS+2% FBS and loaded into SepMate-15™ tubes with 4.5 mL of Lymphoprep™ with a density of 1.077g/mL (StemCell Technologies). Cells were centrifuged at 1200x g for 20min at 4°C. The top layer was collected and centrifuged at 300xg for 8 min. Supernatant was removed and cells were resuspended in PBS + 2% FBS and this step was repeated. Subsequently, red blood cell lysis was performed using the Red Blood Cell Lysis Solution (Miltenyi Biotec) following the manufacturer’s protocol that includes the washing step.

### Single-cell library construction

Single-cell RNA-seq protocol was performed using the Chromium™ Single Cell 3′ reagent kit v2 chemistry and cells were loaded on a GemCode Single Cell Instrument (10x Genomics, Pleasanton, CA). Approximately 5000 single cells were targeted from each sample. Libraries were sequenced on Illumina HiSeq 2500 with the custom configuration of read 1 (26bp) and read 2 (98bp), i7 index (8bp) and i5 index (0bp).

### Single-cell RNA-seq alignment, UMI counting and analysis

The Cell Ranger Single Cell Software Suite 2.1.0 was used to perform single cells demultiplexing and UMI counting (https://support.10xgenomics.com/single-cell-gene-expression/software/overview/welcome). The transcriptome reference used was mm10. Subsequently, the duplicates from young and old mice, respectively, were aggregated using Cellranger aggr. Cells that have more than 500 genes detected and less than 10% of mitochondrial reads were included in the downstream analysis using Seurat 2.3.0. Young and old UMI counts were merged using the MergeSeurat function. Default parameters of Seurat was used in the analysis, unless otherwise stated. 1080 highly variable genes were used as an input for PCA. T-SNE projection and clustering analysis (dims=1:30 and resolution=0.4) were performed using Seurat. Markers genes for each cluster were found using the FindConservedMarkers function and the changes with age within each cell type were identified using the FindMarkers function.

### Statistical analysis

All Student’s t-tests were performed in R and p-values were adjusted using the Benjamini-Hochberg method. Barplots are represented as means with SEM. The top and bottom bounds of boxplots correspond to the 75 and 25th percentile, respectively.

### Data Availability

Single-cell RNA-seq of old and young peripheral blood duplicates are accessible through GEO with the accession number of GSE120505.

## Results

### scRNA-seq of young and old blood identifies different cell types

We performed scRNA-seq using 10x Chromium on peripheral blood obtained from 2 young and 2 old mice. We removed cells that have more than 10% mitochondrial reads, less than 500 genes or more than 4000 genes. 14588 cells that passed this filter (4642 cells from old mouse 1, 2187 cells from old mouse 2, 3902 cells from young mouse 1 and 3857 cells from young mouse 2) were subsequently processed using Seurat/2.3.0 and 17 clusters were obtained (Figure 1a,b). The cell type of each cluster was identified by general marker genes, including Cd3e for T cells, Cd79a and Ms4a2 for B cells, Nkg7 for NK cells, Ly6c2 and Cx3cr1 for monocytes or dendritic cells (DC), Ppbp, Pf4 and Gng11 for megakaryocytes, C1qa, C1qb and C1qc for macrophage, Fcer1a and Cd200r3 for basophil and Hba-a1 for erythrocytes (RBC) (Figure 1c). We identified 5 clusters of T cells, 4 clusters of B cells, 1 cluster of proliferating B or T cells, NK cells, monocytes, dendritic cells, megakaryocytes, macrophage, basophil and (RBC) respectively (Figure 1c).

**Figure 1.**
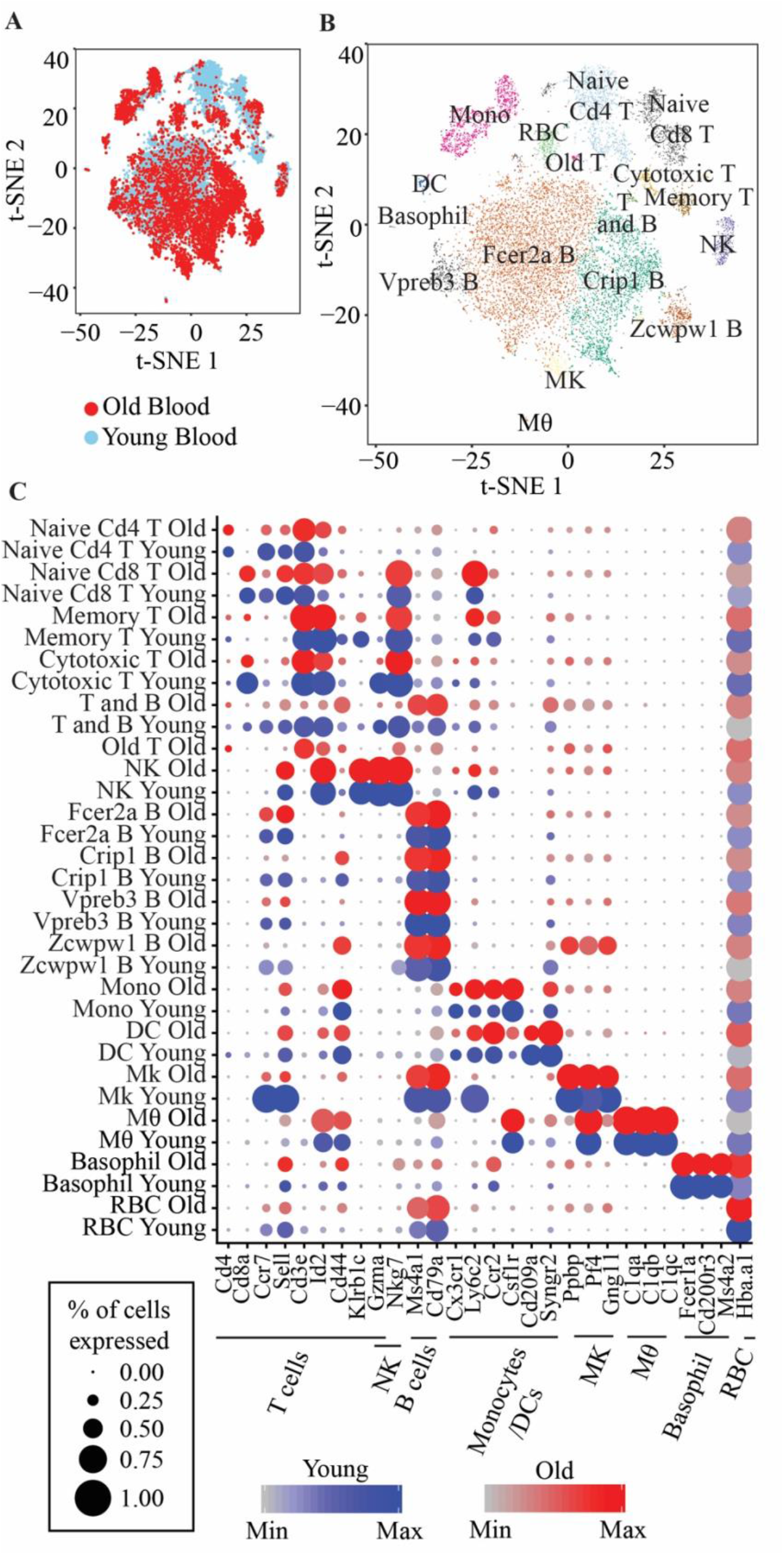
**(A)** t-SNE visualization of 14,588 old and young peripheral blood cells **(B)** t-SNE visualization of the 17 clusters of peripheral blood cells. Memory T: Short-lived effector memory T cells; T and B: Proliferating T and B cells; NK: Natural killer cells; Fcer2a B: Fcer2a, Sell, Ccr7 B cells; Crip1 B: Crip1, S100a6 B cells; Vpreb3 B: Vpreb3, Spib B cells; Zcwpw1 B: Zcwpw1, S100a6 B cells; Mono: Classical monocytes; DC: Monocyte-derived DC; Mk: Megakaryocytes; Mθ: Macrophage; RBC: Red blood cells. **(C)** Marker genes for each immune cell types and the corresponding clusters

The naive T cells cluster is identified by two markers, Sell+ and Cd44-, the memory T cells cluster is identified by Klrb1c+, Id2^high^ and Cd44+ and cytotoxic T cells are identified by high expression of Gzma (Figure 1c). We also identified four clusters of B cells, with one cluster expressing Fcer2a, Sell and Ccr7 marker genes, that we herein refer as Fcer2a B cells, one cluster highly expressing Vpreb3 and Spib genes that we refer to as Vpreb3 B cells, one cluster with Crip1 and S100a6 high expression (Crip1 B cells) and lastly, one cluster of B cells highly expressing Zcwpw1, Lgals1 and Adm (Zcwpw1 B cells) (Supplementary Figure 1b). Classical monocytes are identified by a combination of markers, Ly6c2+, Ccr2+, Sell+ and Csf1r+ whereas monocyte-derived DC are Ly6c2+, Ccr2+, Cd209+ and Syngr2+ (Figure 1c).

### Transcriptomic changes of the same cell type with age

We used Seurat to identify differential gene expression within clusters with age. Only genes that are expressed in at least 10% of cells in each cluster and age group were analyzed. We observed a general trend of the upregulation of genes involved in “ Antigen processing and presentation” and “ Chemokine signaling” pathways with age in naïve Cd4 and Cd8 T cells and Vpreb3 B cells. NK cells from old mice also exhibit increased expression of genes in “ Chemokine signaling” pathway (Figure 2a). In addition, we also observed an upregulation of genes in “ Antigen processing and presentation” in macrophages, Crip1 B cells and Fcer2aB cells. Crip1 B cells and monocytes also showed an upregulation of genes in “ Oxidative phosphorylation” pathway. In contrast, we observed a trend of downregulated genes involved in “ Cytoplasmic Ribosomal Proteins” and “ Ribosome” pathways in several cell types, naïve Cd4 and Cd8 T cells, Crip1 B cells and monocytes, with age (Figure 2a).

**Figure 2.**
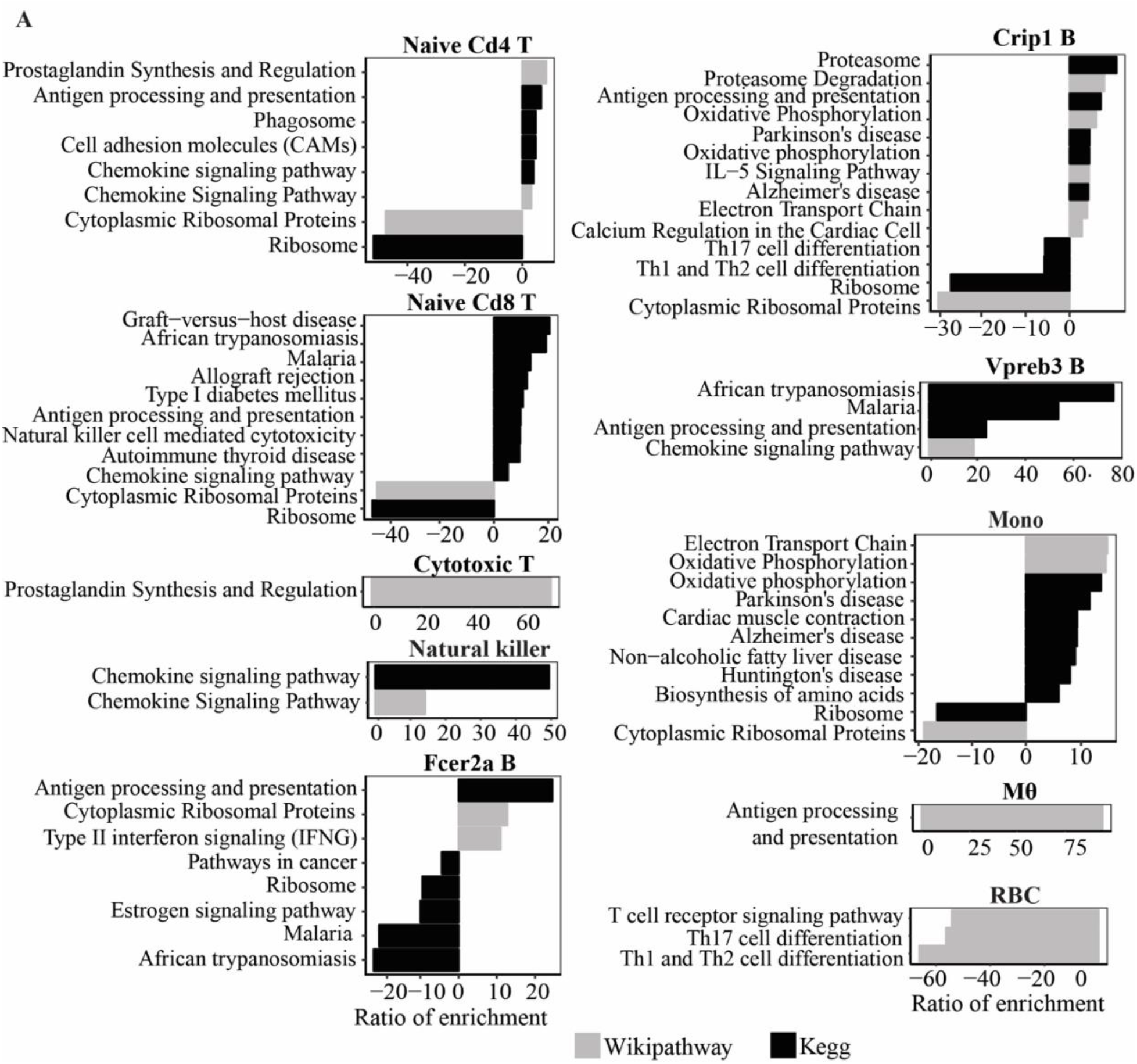
**(A)** Wikipathway and Kegg pathway analysis of upregulated and downregulated genes in old compared to young mice (P<0.05) in each cluster.

### Cell type composition with age

B cells made up the largest proportion of immune cells isolated from the peripheral blood (57.4% in young mice and 70.9% in old mice) (Figure 3a). In young mice, 30.38% of the isolated cells are T cells, 4.66% are monocytes, 4.16% are NK cells, 0.6% are DC, 0.39% are basophils, 0.27% are macrophages and 0.01% are Mk cells. In old mice, 8.41% of the cells are T cells, 10.84% are monocytes, 4.03% are Mk cells, 1.34% are NK cells, 1.09% are DC, 0.44% are macrophages and 0.26% are basophils. Although red blood cell lysis was performed on the samples, we still retrieved 2.06% of RBC from the young mice and 2.68% from old mice. By using the same technique in extracting the cells from young and old mice, we observed a few changes to the composition of cell types with age. First, we detected significantly lower percentage of T cells and NK cells, and higher percentage of B cells and Mk cells in old mice compared to young mice (Figure 3b). Second, within the subsets of B cells, we observed a decrease in the percentage of Fcer2a B cells and increased Crip1 B cells in old mice (Figure 3c). We did not detect any significant changes to the subsets of T cells (Figure 3d).

**Figure 3.**
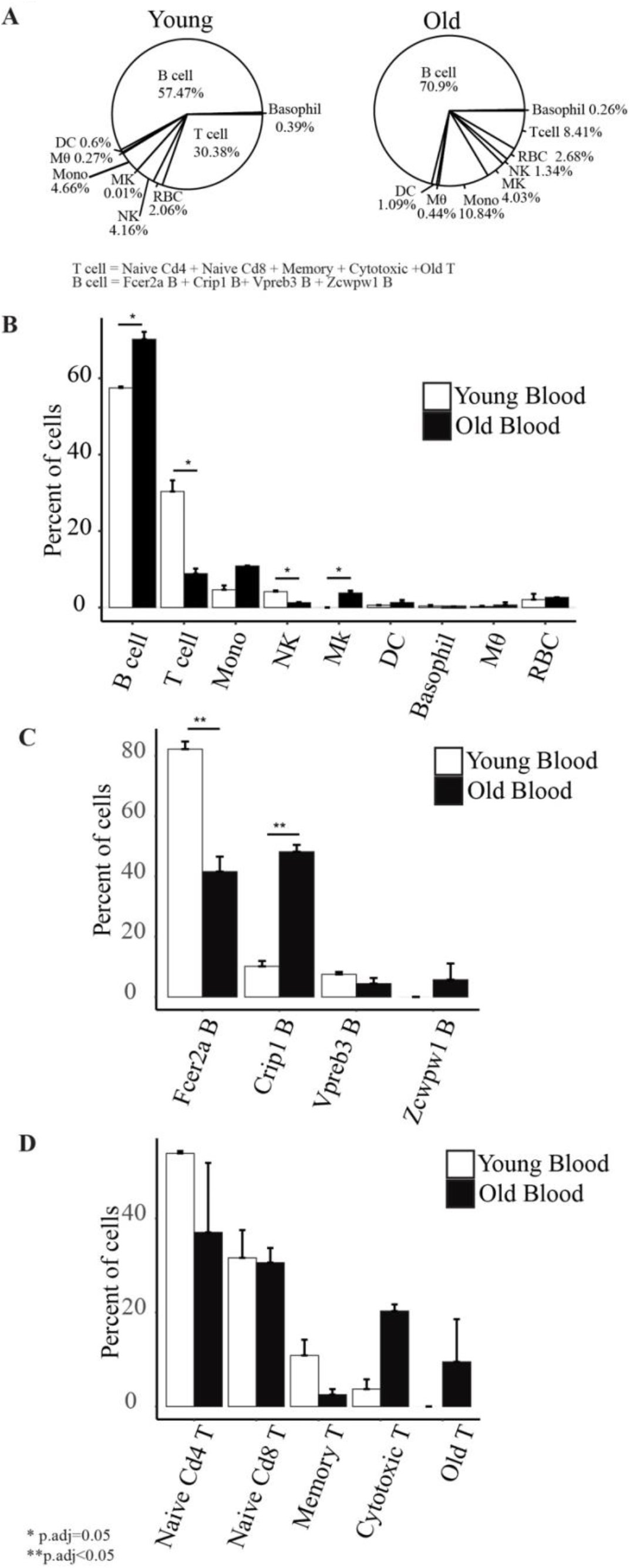
**(A)** Cell type composition of old and young peripheral blood. **(B)** Barplots showing the comparison of the percentage of each cell type with age. **(C)** Barplots showing the comparison of the subset of B cells’ percentage with age. **(D)** Barplots showing the comparison of the subsets of T cells’ percentage with age.

### Immunosenescence

We identified one cluster that consists of only cells from old mice (labeled as “ Old T” in Figure 1b) and these cells expressed Cd3e, the marker of T cells. In addition, this cluster also expressed Cd40lg and Tnfsf8 compared to other T cells clusters, suggesting that it consists of activated T cells (Supplementary Figure 2). A pathway analysis of genes significantly upregulated between this cluster and other T cells clusters showed an enrichment of “ Antigen processing and presentation” and “ NF-kappa B signaling pathway” (Figure 4a). In addition, this cluster expressed significantly higher Bcl2 expression (p=3.9×10^−15^)(Figure 4b) than other T cells clusters. Notably, it also showed significantly higher expression of Cdkn2a and lower expression of Cd28 compared to all other clusters (p=5.38×10^−40^ and p=6.26×10^−8^, respectively) (Figure 4c, Supplementary Figure 2).

**Figure 4.**
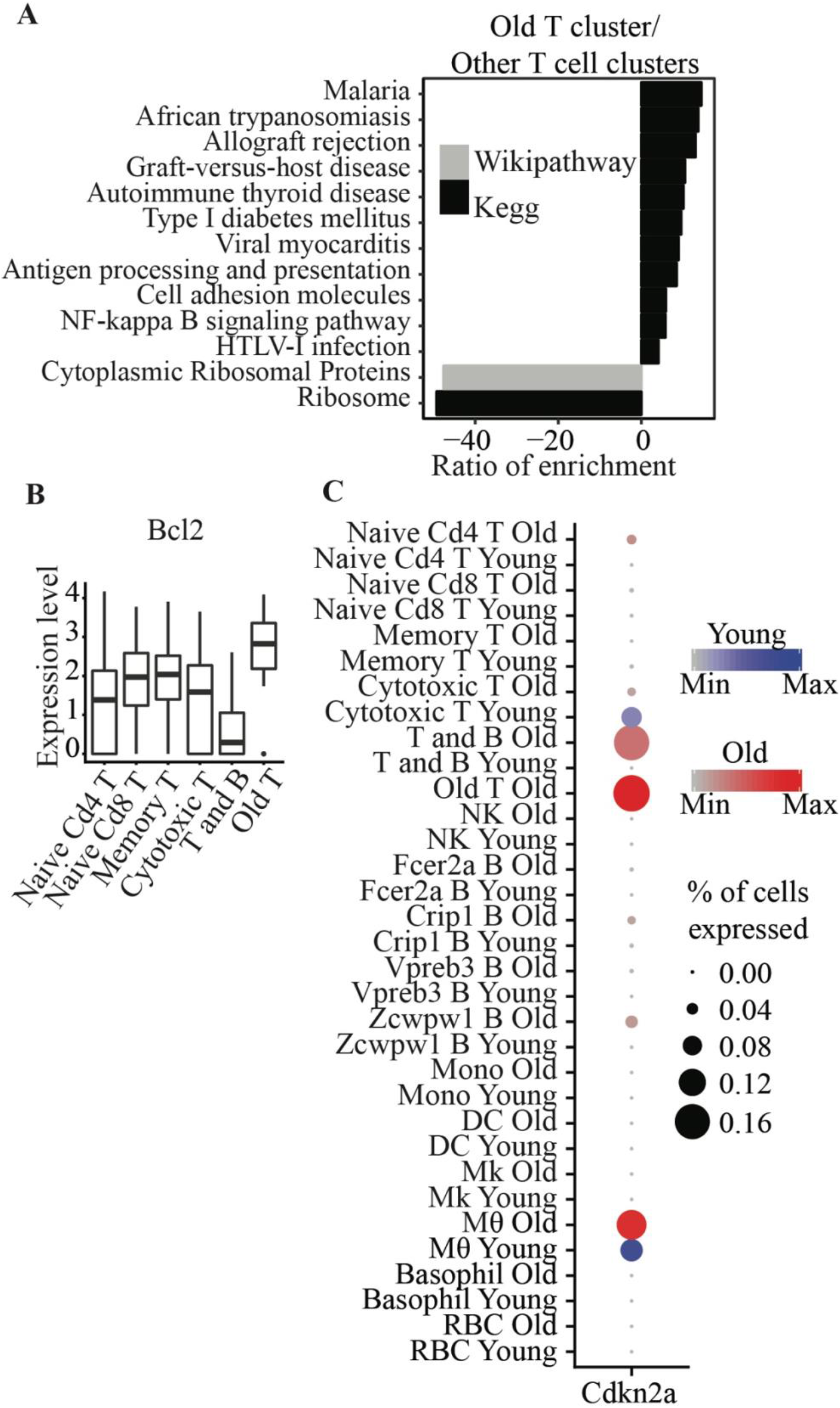
**(A)** Wikipathway and Kegg pathway analysis of differentially expressed genes between Old T cluster and other T cell clusters. **(B)** Bcl2 is significantly higher in Old T cluster compared to other T cell clusters. **(C)** Cdkn2a is significantly higher in Old T cluster compared to all other clusters.

Immunosenescence has been widely implicated in aging. Cellular senescence, an irreversible cell cycle arrest phenomenon was first discovered in fibroblast and senescent cells can be identified by a few markers, including an upregulation of Cdkn1a and Cdkn2a expression. We observed a significant increase of the percentage of old blood cells (6.3%) expressing Cdkn1a and/or Cdkn2a compared to young cells (Fisher’s Exact Test p<2.2×10^−16^) (Figure 5a). However, we did not detect any cluster with significant expression of these two genes between age, but we did observe a moderate (not significant; p=0.066) increase of Cdkn1a expressing cells in naïve Cd4 T cells (Figure 5b).

**Figure 5.**
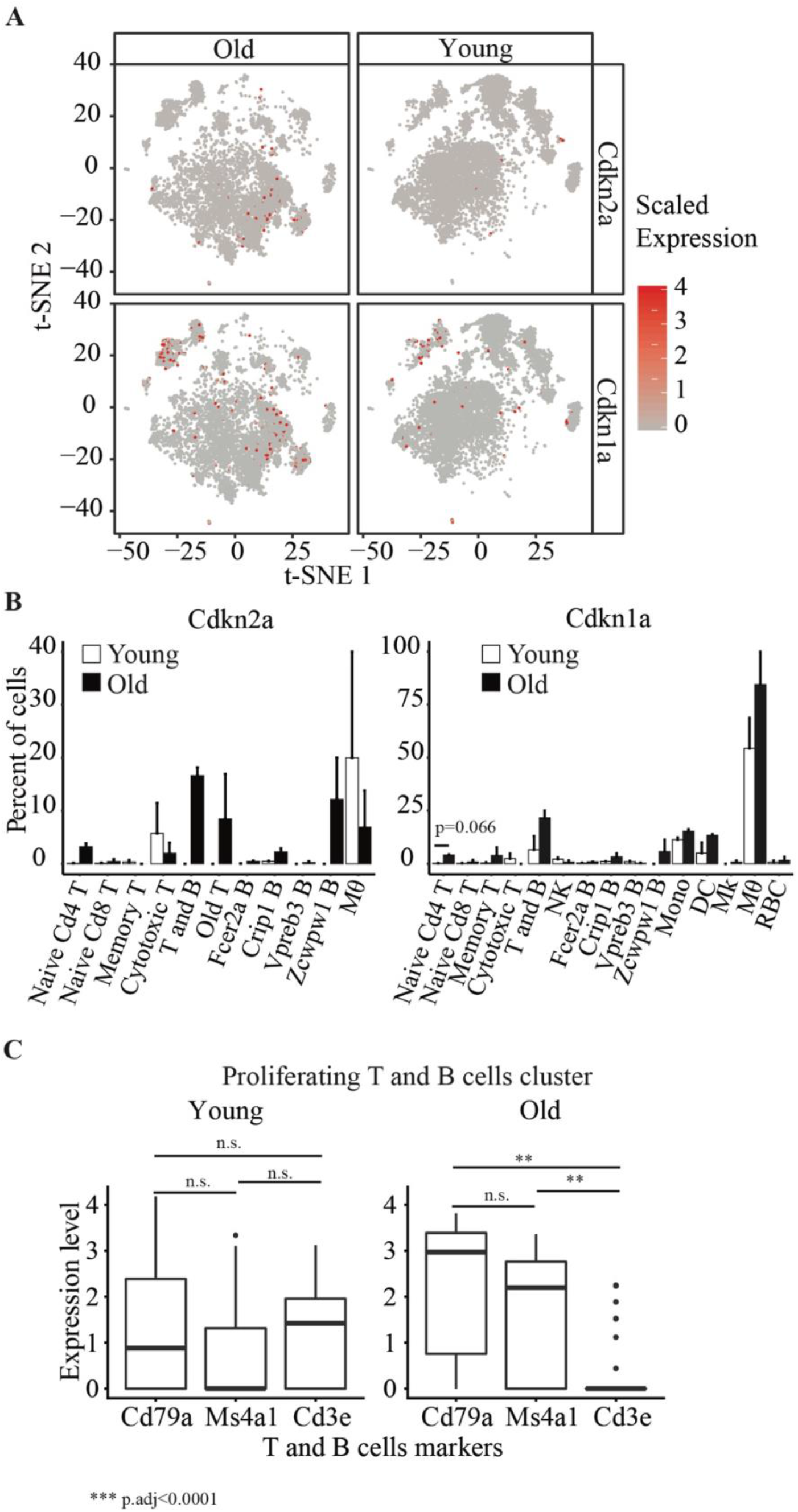
**(A)** t-SNE visualization of cells with different Cdkn2a and Cdkn1a expression levels in young and old. **(B)** Barplots showing percent of cells in young and old expression Cdkn2a and Cdkn1a. (C) Boxplots showing the expression levels of Cd79a, Ms4a1 and Cd3e in the proliferating T and B cells cluster.

We also detected a cluster of old and young proliferating cells that consists of T and B cells (Figure 5C). The cells from old mice in this cluster are enriched in B cells but not T cells markers, whereas we saw a similar enrichment of T and B cells in young mice. This suggests that there is decreased number of proliferating T cells in the old mice.

## Discussion

Transcriptional changes with age in specific immune cell types or lymphoblastoid cell lines in human have been investigated in previous studies (16, 17). scRNA-seq is a powerful technique that can be used to dissect the transcription profiles of thousands or more of cells from the same sample (18). Other studies have used single-cell RNA-seq on the peripheral blood to identify and reclassify cell types and to investigate the changes of immune function of certain cell types upon stimulation with age in human and mice, respectively (15, 19, 20). Here, we profiled 14588 single cells from the mouse blood and assessed the differences between young and old mice.

We identified 17 clusters of cells that we further assigned into different cell types. Immune cells are typically identified using flow cytometry through cell surface markers. (21). We are able to assign various types of T cells but not B cells using markers identified from available studies. Further work is needed to link cell surface markers, immune function and marker genes from the transcriptome of subset of B cells in the peripheral blood.

We found a T cell cluster that is specific to only old mice and these cells have significantly higher Bcl2 expression than all other T cell clusters. Bcl2 is an anti-apoptotic factor that regulates cell death and it has been used as a target for senolytic drugs to clear senescent cells. This cluster also exhibit significantly higher expression of Cdkn1a, which is a senescent marker and an anti-proliferative marker that have been found in T cells (22, 23). Senescent T cells have been previously observed in not only aged samples, but also patients with cancer or autoimmune disease (24). A previous study showed that T cells that are antigen-induced to cell death in vitro can be rescued by the expression of p16 (25). Another study showed that p16 reduces naïve T cell and memory T cell proliferation and the deletion of p16 in T cell lineage attenuates aging phenotypes associated with T cells (26). Among other T cells, memory T cells are known to exhibit higher expression of Bcl2 (27). Loss of Cd28 has also been a reported in replicative senescent cells (28). Notably, we only detected this in old mice. It has been previously suggested that an accumulation of CD28-T cells in older individuals may be due to increased proliferation instead of the cells being more resistant to apoptosis with age (29, 30). However, the higher expression of Cdkn1a and Bcl2 genes and a lower expression of Cd28 in the old T cluster in our dataset may indicate that these cells are senescent and more resistant to apoptosis than other T cells.

In general, we observed an enrichment of chemokine signaling and antigen presenting pathways with age. This may indicate that the immune system works at a higher capacity in old mice. There is also increased number of cells with Cdkn2a and Cdkn1a expression in old age but the increased is not specific to any cell type, suggesting that there is a general increased of possibly senescent cells.

The changes in the cell type composition that we observed is consistent with previous studies, such as increased B cells in old peripheral blood in mice, decreased number of NK cells and T cells in aged mouse blood (3, 5, 31). In addition, we also observed an increased in Mk cells with age. To our knowledge, the change of the number of this cell type has not been documented before, but platelet counts were shown to be relatively stable until old age, where it starts decreasing in human. On the other hand, platelet count was shown to increase in 18 months old mice compared to young mice (32) but another study showed that the count does not change in 24-25 months mice compared to young (33).

Finally, targeting senescent cells using genetic approaches have been shown to ameliorate aging phenotype (34, 35). More recently, senolytics drugs are being identified or developed to target apoptotic pathways because senescent cells are known to be apoptosis-resistant (34). Therefore, the Bcl2+ old T cells that we identified in old mice can potentially be targeted pharmacologically to ameliorate the phenotypes associated with the aging of the immune system.

## Acknowledgement

Part of this research was conducted using computational resources at the Center for Computation and Visualization, Brown University.

---

**Supplementary Figure 1.**
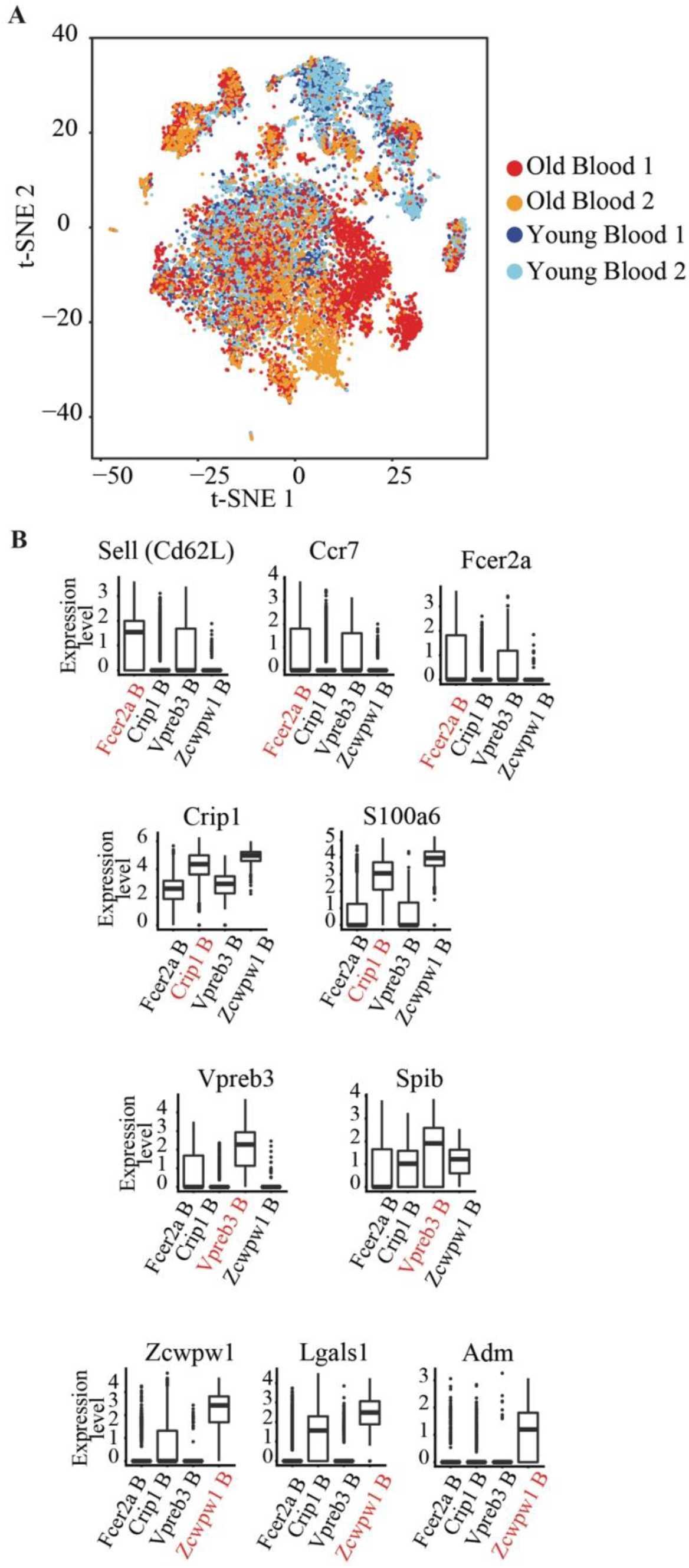
**(A)** t-SNE visualization of young and old blood replicates. **(B)** Boxplots showing the expression levels of the respective marker genes in each B cells clusters. The cluster that the B cells are identified in were indicated in red.

**Supplementary Figure 2.**
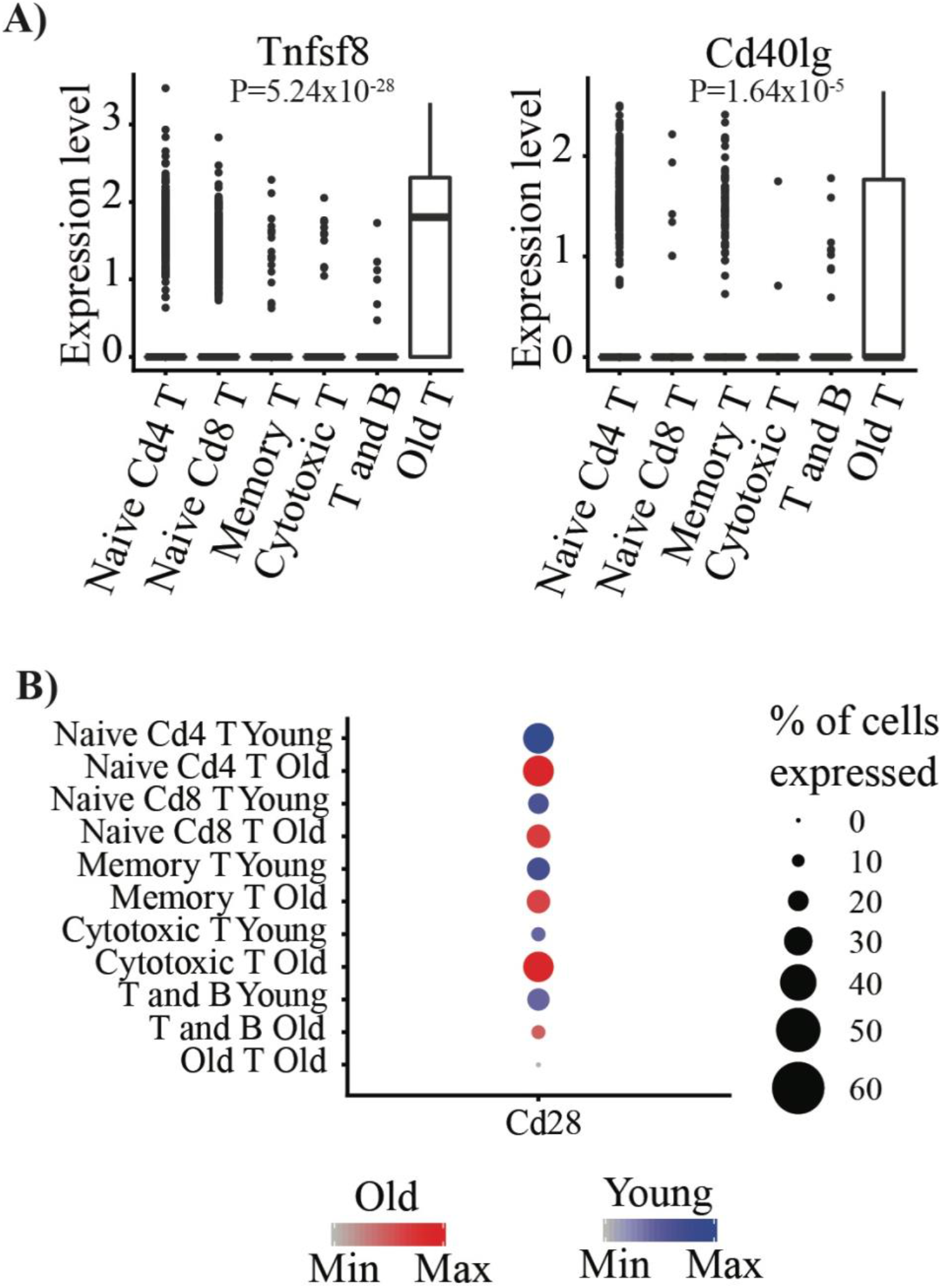
**(A)** Expression level of Tnfsf8 and Cd40lg in T cells clusters. **(B)** Expression level of Cd28 in Old T cluster and other T cell clusters.

